# Enhancing Prime Editing Efficiency and Flexibility with Tethered and Split pegRNAs

**DOI:** 10.1101/2022.04.05.487236

**Authors:** Ying Feng, Siyuan Liu, Qiqin Mo, Xiao Xiao, Pengpeng Liu, Hanhui Ma

**Affiliations:** Gene Editing Center, School of Life Science and Technology, ShanghaiTech University, Shanghai; China & School of Biotechnology, East China University of Science and Technology, Shanghai, China; Gene Editing Center, School of Life Science and Technology, ShanghaiTech University, Shanghai; School of Biotechnology, East China University of Science and Technology, Shanghai, China; Department of Molecular, Cell and Cancer Biology, University of Massachusetts Medical School, Worcester, MA, USA

**Author notes:** Correspondence should be addressed to Hanhui Ma. These authors contributed equally (Ying Feng, Siyuan Liu).

## Abstract

Prime editing (PE) has advantages for small insertion, deletion or point mutations without double-stranded DNA breaks. The 3’-extension of pegRNAs could negatively affect its stability or folding and comprise the PE activity. Here we generated <u>s</u>tem-loop PEs (sPEs) by adding stem-loop aptamers at the 3’-terminal of pegRNA, which can be tethered to Cas9 nickase resulting in tethered PEs (tPEs). sPEs and tPEs increased the small insertion, deletion or point mutations efficiency by 2-4-fold on average in HEK293, U2OS and HeLa cells. We split the modified pegRNAs into sgRNA and prime RNA. The resulting <u>s</u>plit pegR<u>N</u>A prime editors (SnPEs) maintain the PE activity and increase flexibility.

## Introduction

Most human genetic diseases arise from mutations such as insertion, deletion or point mutations (Lamdrum et al., 2016). CRISPR-Cas system has been repurposed to correct pathogenic mutations in a variety of genetic diseases (Cong et al., 2013; Mali et al., 2013). There are many concerns about using CRISPR-mediated double-stranded DNA breaks (DSBs) for therapeutic purposes, primarily due to off-targeted mutations (Kosicki et al., 2018). Base editing can efficiently modify single nucleotide mutations without DSBs in dividing and post-mitotic cells (Komor et al., 2016; Gaudelli et al., 2017; Kurt IC et al., 2021; Zhao D et al., 2021; Koblan et al., 2021). Nevertheless, base editing can’t correct deletions, insertions, or some point mutations such as transversion mutations (Gaudelli et al., 2017). Prime editing has its advantages of precisely correct point mutations, small insertions or deletions in animal cells (Anzalone et al., 2019) and plants (Lin et al., 2020). However, prime editing efficiency varies among genomic sites or cell types (Nelson et al., 2021; Chen et al., 2021). The reasons for cause variable efficiency of the prime editing is yet to be identified. Prime editing requires the assembly of the prime editor (Cas9 nickase fused to reverse transcriptase) and pegRNA to be PE-pegRNA complex. PE-pegRNA complex searches and nicks target DNA at the non-template strand, followed by reverse transcription, and mutagenesis is done by 3’-flap resolution (Anzalone et al., 2019). Thus, it is crucial to optimize pegRNA and PE-pegRNA complex for higher PE efficiency and precision.

## Results

Robust prime editing is required to satisfy a series of conditions, such as stable and properly folded pegRNAs, effective assembly of PE-pegRNA complex, targeting to genomic loci, efficient reverse transcription and correct editing. Unstructured RNA sequence appended to the 3’-end of sgRNA destabilizes the sgRNAs (Nelson et al., 2021; Zalatan et al., 2015). The prime editors consist of a *Streptococcus pyogenes* Cas9 nickase-H840A with C-terminal fusion of an MMLV (PE2), and a pegRNA which includes a prime binding site (PBS) and a reverse transcription template (RTT) at 3’-terminal of sgRNA. PBS and RTT at the 3’-terminal of pegRNA are easy to be partially degraded, resulting in truncated pegRNAs. The truncated pegRNAs can still search and recognize the target sites, but not be able to complete the correct editing due to loss of the PBS or RTT-PBS (Nelson et al., 2021). In addition, pegRNA circularization might also result in self-inhibition and compromise the PE efficiency (Liu et al., 2021). We have shown that the dynamics of CRISPR DNA targeting limits genome editing efficiency (Ma et al., 2016a).

Here we used CRISPR-based genome imaging (Ma et al., 2016b; Ma et al., 2018) to compare the target efficiency of CRISPR-based GE (Genome Editor) and PE (Prime Editor). Fluorescent Cas9-sgRNA complex effectively targeted to chromosome 3-specific tandem repeats (C3) allows to be visualized under microscopy in U2OS cells (**Figure S1**) (Ma et al., 2016a). As shown in **Figure S1B and S1C**, 2-4 bright foci were observed in the GE system but not the PE system suggesting that 3’-terminal RTT-PBS of pegRNA resulted in low target efficiency of PE. We added the stem-loop aptamer MS2 (Convery et al., 1998) at the 3’-terminal of pegRNAs (pegRNA-MS2) and found that visualization of C3 loci was recovered (**Figure S1B-S1C**). We assume that the 3’-terminal pegRNA tethered to Cas9 nickase will stabilize the PE-pegRNA complex. We fused tandem MS2 coat protein (tdMCP) to the N-terminal of Cas9 for binding 3’-terminal MS2 at the engineered pegRNA. As we can see in **Figure S1B-S1C**, C3 labeling was maintained. The low targeting efficiency of canonical PE suggests that inefficient targeting of genomic loci may compromise the PE efficiency. On the contrary, the recovery of C3 loci visualization in 3’-terminal MS2 tagged pegRNA or tethered to Cas9 nickase indicates the engineered pegRNA or tethered to Cas9 nickase may improve the PE efficiency.

To distinguish from the canonical prime editor (PE), we named the PE system with 3’-stem-loop MS2, PP7, Csy4 and BoxB tagged pegRNA to be stem-loop PE (sPE), and the system with pegRNA-MS2, PP7, Csy4 and BoxB tethered to Cas9 nickase-MMLV was named tethered PE (tPE) (**Figure 1A**). First, we tested sPE-MS2, PP7, Csy4 and BoxB and tPE-MS2, PP7, Csy4 and BoxB (Urbanek et al., 2014; Haurwitz et al., 2010) on the PE efficiency using PE3 in HEK293FT cells at *RUNX1* (+5 G·C to T·A). Use of either sPEs or tPEs improved correct editing efficiency with no significant change in edit/indel ratios (**Figure 1B, S4A, S4D**). We also compared ePE-Mpknot, ePE-EvopreQ1 (Nelson, J.W et al., 2021) on the same loci in HEK293FT cells. The correct editing efficiency of sPE-MS2, PP7, Csy4, BoxB, and tPE-MS2, Csy4 on this loci are higher than ePE-Mpknot, EvopreQ1 (**Figure 1B**). We also tested the small insertion and deletion efficiency by tPE-MS2, PP7, Csy4 and BoxB at *RUNX1*_with +1 ATG insertion (**Figure 1C**) or_+1 CGA deletion (**Figure 1D**) resulting in a 4.9 or 2.7-fold increase on average in PE efficiency with no significant change in edit/indel ratios overall (**Figure S4B-4C, S4E-4F**) relative to that of canonical PE in HEK293FT cells. Therefore, we chose MS2 appended at the 3’-terminal of pegRNA on the PE efficiency using PE3 in HEK293FT cell at ten loci including *SRD5A3* (+2 C·G to A·T), *DYRK1A* (+1 C·G to G·C), *HDAC1* (+1 C·G to G·C), *BCL11A* (+1 C·G to A·T), *GFAP* (+1 A·T to T·A), *RUNX1* (+5 G·C to T·A), *JAK2 (*+1 C·G to T·A), *SRD5A1* (+1 C·G to A·T), *DMD (*+1 T·A to C·G) and *EED* (+1 A·T to T·A) (**Figure 1E-1F**). Use of either sPE-MS2 or tPE-MS2 resulted in a 1.8 or 1.9-fold average improvement in PE efficiency relative to that of canonical PE across tested sites in HEK293FT cells (**Figure 1G**) with no significant change in edit/indel ratios overall (**Figure S3A-3B**). We also test the insertion efficiency of *DNMT1*_+4 A, +4 AC and +4 ACT and deletion efficiency of *DNMT1*_+4 G, +4 GG and +4 GGG using tPE-MS2. The PE efficiency by tPE-MS2 increased 2-3 folds on average without significant change in edit/indel ratios (**Figure S5**).

**Figure 1.**
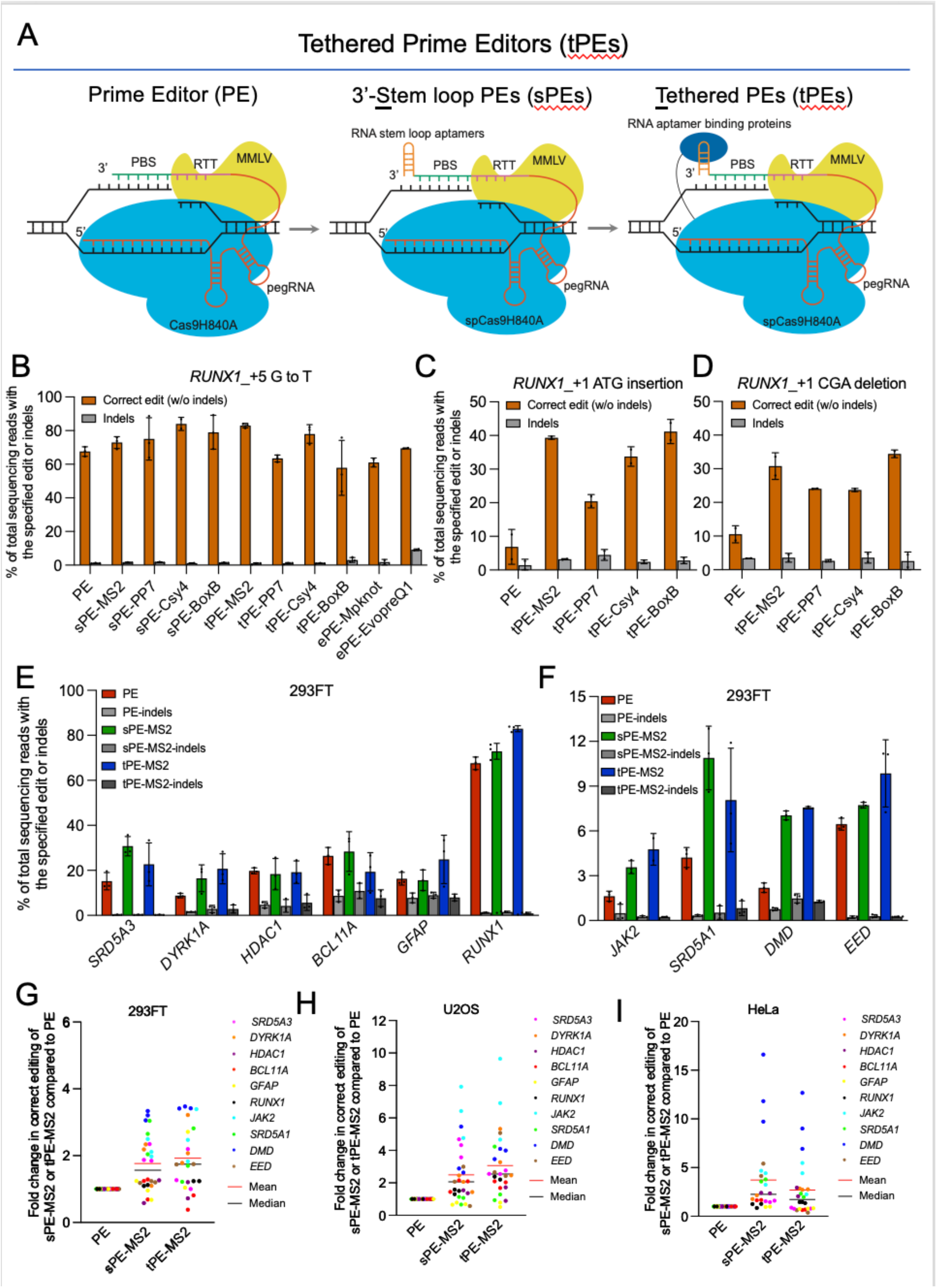
pegRNA with 3’-RNA aptamers or tethered to Cas9 nickase enhance targeting and editing efficiency. (A) The prime editing (PE) complex consists of a *Streptococcus pyogenes* Cas9 nickase-H840A (sky blue) with C-terminal fusion of an MMLV (yellow), and a pegRNA which includes a prime binding site (PBS, bluish-green) and a reverse transcription template (RTT, reddish-purple) at 3’-terminal of sgRNA. 3’-stem-loop PE (sPE)-MS2 was generated by appending an MS2 stem-loop aptamer (orange) to the 3’-terminal of pegRNA. The tethered PE (tPE)-MS2 was generated by fusing tandem MS2 coat protein (tdMCP, blue) to the N-terminal of Cas9 for cognate RNA aptamers in sPEs. (B) comparison of editing efficiency between PE, sPE-MS2, PP7, Csy4, BoxB, tPE-MS2, PP7, Csy4, BoxB, ePE-Mpknot, EvopreQ1 mediated point mutation of *RUNX1*_+5 G·C to T·A using PE3 in HEK293FT cells. (C) Editing efficiency for *RUNX1*_+1 ATG insertion of tPE-MS2, PP7, Csy4 and boxB. (D) Editing efficiency for *RUNX1*_+1 CGA deletion of tPE-MS2, PP7, Csy4 and boxB, compared to canonical PE (dashed line). (E) The efficiency of PE, sPE-MS2 and tPE-MS2 mediated point mutation of *SRD5A3_*+2 C·G to A·T, *DYRK1A_*+1 C·G to G·C, *HDAC1_*+1 C·G to G·C, *BCL11A*_+1C·G to A·T, *GFAP*_+1A·T to T·A and *RUNX1*_+5 G·C to T·A using PE3 in HEK293FT cells. (F) The efficiency of PE, sPE-MS2 and tPE-MS2 mediated point mutation *JAK2 _*+1 C·G to T·A, *SRD5A1* _+1 C·G to A·T, *DMD_*+1 T·A to C·G and *EED_* +1 A·T to T·A using PE3 in HEK293FT cells. Comparison of editing efficiencies of canonical PE, sPE-MS2 or tPE-MS2 for point mutation at ten loci in HEK293FT cells (G), U2OS (H) and HeLa cells (I). Values were calculated from the data presented in Figure 1 and Figure S2. Dots indicate the average of three biological replicates and Bars indicate the grand median. Data and error bars in (B-F) indicate the mean and standard deviation of three independent biological replicates.

PE efficiency varies in different cell types (Nelson et al., 2021; Chen et al., 2021). To ensure that the improvement in PE efficiency by sPEs or tPEs was not limited to HEK293FT cells, we tested the above ten loci with sPE-MS2 or tPE-MS2 in U2OS (**Figure S2A-2B**) and HeLa cells using PE3 (**Figure S2C-2D**). In either U2OS or HeLa, sPE-MS2 or tPE-MS2 resulted in improvements in editing efficiency compared to canonical PE, averaging 2.5 or 3.1-fold higher editing in U2OS cells (**Figure 1H**) and 3.7 or 2.7-fold higher editing in HeLa cells (**Figure 1I**), with no significant change in edit:indel ratios overall (**Figure S3A-3D**). These results indicate that sPE and tPE can enhance PE efficiency in different cell types. We examined off-target editing by sPE-MS2 and tPE-MS2 for *DYRK1A, RUNX1, BCL11A, SRD5A3* and *JAK2* loci in HEK293FT cells. The off-target sites were predicted by Cas-OFFinder (Bae et al., 2014). Average <0.1% off-target prime editing was detected in canonical PE, sPE-MS2 and tPE-MS2 at the predicted off-target sites for each protospacer of *DYRK1A, RUNX1, BCL11A, SRD5A3* or *JAK2* in HEK293FT cells (**Figure S6**).

To make the PE more flexible, we split the modified pegRNA in the tPEs into sgRNA and <u>p</u>rime RNA (pRNA), and generate <u>s</u>plit pegR<u>N</u>A prime editors (SnPEs) (**Figure 2A**). To stabilize pRNA, we generate <u>c</u>ircular <u>p</u>rime RNA (cpRNA) by Tornado circRNA expression system (Litke et al., 2019) (**Figure S7**). Briefly, we split pegRNA-MS2, PP7, BoxB and Cys4 into sgRNA and pRNA, resulting in pRNA-5’-MS2, pRNA-3’-MS2, pRNA-<u>c(</u>ircular)-MS2, pRNA-5’-PP7, pRNA-3’-PP7, pRNA-c-PP7, pRNA-5’-BoxB, pRNA-5’-Cys4 (**Figure 2A**). Very low SnPE activity was observed when using control pRNA without MS2 or PP7 in U2OS, HEK293FT and HeLa cells (**Figure 2B-2E**). The PE efficiency is comparable to canonical PE when SnPE-5’-MS2, SnPE-c-MS2, SnPE-5’-PP7 were used. It shows 82.3% of canonical PE activity for SnPE-5’-MS2, 72.1% for SnPE-c-MS2 and 59.4% for SnPE-5’-PP7 and 46.0% for SnPE-c-PP7 in HEK293FT cells (**Figure 2B-2E**). The efficiency became much lower when SnPE-3’-MS2 (31.7%) or SnPE-3’-PP7 (18.3%) was used (**Figure 2B-2E**). The highest PE efficiency was found when using SnPE-5’-PP7 (79.5% of canonical PE) in U2OS and SnPE-5’-MS2 (59.8%) in HeLa cells. The edit/indel ratios of SnPEs are slightly lower than canonical PE in all three cell types, which is in concord with the level changes of PE activities (**Figure S8A-8F**).

**Figure 2.**
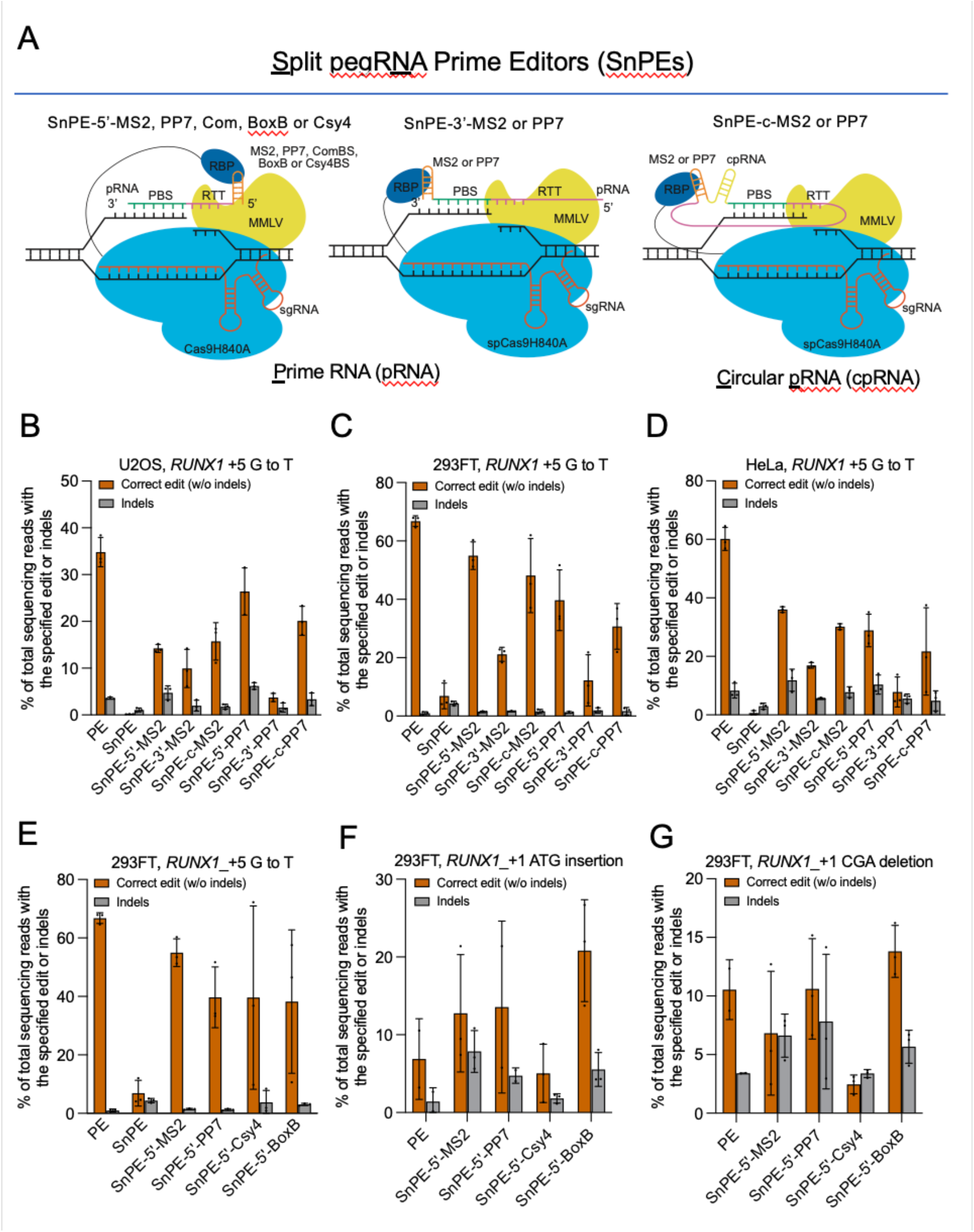
Split pegRNA prime editing maintains PE activity by tethering prime RNA to Cas9. (A) Schematics of split pegRNA prime editors. The <u>s</u>plit pegR<u>N</u>A prime editor (SnPE) consists of a Cas9 nickase-H840A (sky blue) fused with C-terminal MMLV (yellow) and N-terminal RNA binding proteins (RBPs, blue), a sgRNA, and a separated prime RNA (pRNA). pRNA consists of a prime binding site (PBS, bluish-green), a reverse transcription template (RTT, reddish-purple) and a RNA stem-loop aptamer such as MS2 or PP7 (orange). SnPE-5’ or 3’-MS2 or PP7 consists of RBP-Cas9 nickase-MMLV, a pRNA bearing MS2 or PP7 and a sgRNA. RBP at the N-terminal Cas9 binds the RNA aptamer MS2 or PP7. SnPE-c-MS2 or PP7 includes the circular pRNA (cpRNA) bearing MS2 or PP7 generated by the tornado circular RNA expression system. Efficiency of SnPE-5’-MS2 or PP7, SnPE-3’-MS2 or PP7, SnPE-c-MS2 or PP7 were tested at *RUNX1*_+5 G·C to T·A using PE3 in U2OS cells (B), HEK293FT cells (C), and HeLa cells (D). (E) Efficiency of PE, SnPE, SnPE-5’-Com, SnPE-5’-BoxB and SnPE-5’-Csy4 at *RUNX1*_+5 G·C to T·A using PE3 in HEK293FT cells. (F) Efficiency of SnPE-5’-MS2 or PP7, SnPE-5’-MS2 or PP7, SnPE-5’-Csy4 and SnPE-5’-boxB were tested at *RUNX1*_+1 ATG insertion using PE3 HEK293FT cells. (G) Efficiency of SnPE-5’-MS2 or PP7, SnPE-5’-MS2 or PP7, SnPE-5’-Csy4 and SnPE-5’-boxB were tested at *RUNX1*_+1 CGA deletion using PE3 HEK293FT cells. Data and error bars in (B-G) indicate the mean and standard deviation of three independent biological replicates.

We further tested whether pRNA-5’-BoxB or pRNA-5’-Cys4 could also improve the *RUNX1*_+1 ATG insertion efficiency and *RUNX1*_+1 CGA deletion efficiency. The SnPE-5’-MS2, PP7, BoxB showed higher activities for *RUNX1*_+1 ATG insertion than canonical PE in HEK293FT cells, particularly the SnPE-5’-BoxB showed two folds increase in the efficiency for *RUNX1*_+1 ATG insertion (**Figure 2F**). SnPE-5’-PP7, BoxB also showed higher activities for *RUNX1*_+1 CGA deletion than canonical PE (**Figure 2G**) in HEK293FT cells. The edit/indel ratios changes are in concord with the level changes of PE activities (**Figure S8G-8L**). These results indicate SnPEs maintain the activity and increase the flexibility of prime editing.

## Discussion

The Prime editing efficiency varies among genomic loci or cell types. For instance, the PE efficiency of *EED* (+1 A·T to T·A) is very lower (0.3% of correct editing), while the PE efficiency of *RUNX1* (+5 G·C to T·A) reaches 80.0% of correct editing in U2OS cells (**Figure 1B**). The efficiency of prime editing in HEK293 shows generally higher than in HeLa and U2OS cells. The Mechanisms of variable PE efficiency at different loci in different cell types are still not clear. Here we applied for CRISPR-based genome imaging to study target recognition and found low targeting efficiency of canonical PE, which can be restored by sPE-MS2 or tPE-MS2. It could be possible that sPE-MS2 or tPE-MS2 stabilizes of pegRNAs, facilitates the assembly of PE2-pegRNA complexes, or promotes target recognition by PE2-pegRNA complex. It will be interesting to examine the target efficiency of other engineered pegRNAs such as epegRNAs (Nelson et al., 2021), different length of PBS or mismatches on RTT-PBS (Anzalone et al., 2019) and optimize the prime editing efficiency.

The prime editing has the advantages for point mutations, small deletions and insertions. However, the instability or misfolding of pegRNAs may have limited its applications for direct insertion of bigger size fragment such as >100 nucleotides. It will be interesting to test whether sPEs or tPEs will allow for the installation of DNA fragments with hundreds of nucleotides. There are several dual pegRNA strategies to increase the efficiency and precision of small or large deletions, small fragment insertions (Choi et al., 2021; Jiang et al., 2021; Anzalone et al., 2021). Large fragment insertion has also been achieved by the combination of dual-pegRNA mediated small insertions and recombinase-mediated site-specific genomic integration (Anzalone et al., 2021). It will be intriguing to test whether sPEs or tPEs will benefit these dual-pegRNA systems since sPEs and tPEs showed better targeting efficiency than canonical PEs.

Tethered PEs offer the opportunity to liberate the RTT-PBS unit from the pegRNAs and spatiotemporally control the PEs. pegRNA in tPEs was separated to be conventional sgRNA and prime RNA containing PBS, RTT and RNA aptamer resulting in SnPEs. One of the potential applications of SnPEs is more readily prepared by chemical synthesis (Hendel et al., 2015; Yin et al., 2017) of split pegRNAs due to the smaller sizes of sgRNA and pRNA. Synthetically modified sgRNA and prime RNAs may further enhance the PE efficiency. The SnPEs could also combine with inhibitors of DNA mismatch repair (MMR) (Chen et al., 2021) to further increase prime editing efficiency and precision. Separated prime RNA could be also introduced under the control of chemicals or lights (Stanton et al., 2018; Zhou et al., 2021) and evolve the PE system to be tunable in space and time.

## Acknowledgements

This work was funded by National Natural Science Foundation of China (No. 31970591 to H. Ma), the Shanghai Pujiang program (19PJ1408000 to H. Ma) and Shanghai Science and Technology Innovation Action Plan (21JC1404800 to H. Ma). 293FT cell was a gift from W. Qi Lab. We thank P.W. Zhang and L.S. Zhang for help with cell sorting and H.Z. Liu for imaging. We are grateful to Biomedical Big Data Platform L.C. Jiang and Z.Y. Song for deep sequencing. Fluorescence activated cell sorting (FACS) and DeltaVision Ultra microscopy were provided by Shanghai Institute for Advanced Immunochemical Studies (SIAIS) at Shanghaitech University.

## Author contributions

H.M. conceived the idea. Y.F. and S.L. designed the research, performed experiments and analyzed data. P.L. and Q.M. analyzed deep-sequencing data. H.M., X.X. and Y.F. wrote the paper.

## Methods

### Plasmid construction

Modified pegRNAs or split pegRNAs expression plasmids pPB-PE-ALL-RNA, pPB-sPE-ALL-RNA, or pPB-SnPE-ALL-RNA for sPEs tPEs or SnPEs were generated by two steps (Figure S9). First, pPE-pegRNA, psPE-pegRNA, pSnPE-5’-pRNA, pSnPE-3’-pRNA, pSnPE-circ-pRNA and pSnPE-sgRNA pPE3-sgRNA were constructed separately. For pPE-pegRNA, psPE-pegRNA, pSnPE-5’-pRNA, pSnPE-3’-pRNA and pSnPE-circ-pRNA (Table S1), the oligos were synthesized and annealed with 5’ ACCG and 3’ AAAA overhangs and cloned into pLH-AAAA backbone plasmid (Sequence S1). For pSnPE-sgRNA and pPE3-sgRNA, the oligos were synthesized and annealed with 5’ ACCG and 3’ AAAC overhangs and cloned into pLH-sgRNA3 backbone vector (Sequence S1). Second, Golden Gate assembly was used to clone related RNA expression cassettes into one vector pDONOR5.1 (sequence S1) and generate pPB-PE-ALL-RNA, pPB-sPE-ALL-RNA, or pPB-SnPE-ALL-RNA respectively. pCMV-PE2 (#addgene 132775) was used to generate RBP-PE2 (SpCas9H840A-MMLV) expression plasmids. First, T2A-GFP was inserted at the C terminal of pCMV-PE2 resulting in pCMV-PE2-T2A-GFP. Second, RNA binding proteins for stem loop aptamers including MCP, tdMCP, PCP, tdPCP, Com, N22p and Csy4H29A (Sequence S2) were synthesized and cloned to the pCMV-PE2-T2A-GFP using ClonExpress II One Step Cloning Kit (Vazyme, C112-01), resulting pCMV-MCP-PE2-T2A-GFP, pCMV-tdMCP-PE2-T2A-GFP, pCMV-PCP-PE2-T2A-GFP, pCMV-tdPCP-PE2-T2A-GFP, pCMV-Com-PE2-T2A-GFP, pCMV-N22p-PE2-T2A-GFP and pCMV-Csy4H29A-PE2-T2A-GFP respectively.

### Cell culture and transfection and genomic DNA extraction

HEK293FT (Thermo Fisher Scientific), HeLa (ATCC, CCL-2), and U2OS (ATCC, HTB-96) cells were cultured in DMEM with high glucose in 10% FBS (fetal bovine serum, Thermo Fisher Scientific). All cells were cultured at 37°C and 5% CO2 in a humidified incubator. Cells were seeded in 12-well plates and transfected at approximately 60% confluence using Lipofectamine 2000 (Thermo Fisher Scientific) according to the manufacturer’s protocols. A total of 1.5 μg Prime editor 2 (PE2, dCas9-MMLV) or RBP-PE2, and 500 ng PE-all-RNA, sPE-all-RNA, tPE-all-RNA or snPE-all-RNA expression plasmids were co-transfected into HEK293FT, U2OS or HeLa cells. 72 h after transfection, Prime editors (GFP+) and PE-all-RNA, sPE-all-RNA, tPE-all-RNA or snPE-all-RNA (BFP+) double-positive cells were collected from flow cytometry (BD FACS Aria III). The genomic DNA of cells was extracted using QuickExtract DNA Extraction Solution (Lucigen) according to the manufacturer’s protocols. The isolated DNA was PCR-amplified with Phanta Max Super-Fidelity DNA Polymerase (Vazyme). Primers used are listed in Table S2.

### High-throughput DNA sequencing of genomic DNA samples

Genomic sites of interest were amplified from genomic DNA using locus-specific primers containing the Truseq adapters. Equal amounts of PCR products were pooled and gel purified. The purified library was deep sequenced using a paired-end 150 bp Illumina MiniSeq run. All prime editing experiments were analyzed as follows. Demultiplexing and base calling were both performed using bcl2fastq Conversion Software v2.18 (Illumina, Inc.), allowing 0 barcode mismatches with a minimum trimmed read length of 75. Alignment of sequencing reads to the Amplicon sequence (Table S3) was performed using CRISPResso2 (Clement et al., 2019) in standard mode using the parameters ‘‘-q 30’’.

For each amplicon, the CRISPResso2 quantification window was positioned to include the entire sequence between pegRNA- and sgRNA-directed Cas9 cut sites, as well as an additional 10 bp beyond both cut sites. For PE activity quantifications from deep sequenced data, editing efficiency was calculated as the percentage of reads with the desired editing without indels (‘‘-discard_indel_reads TRUE.’’ mode) out of the total number of reads ((number of desired editing-containing reads)/(number of reference-aligned reads).). For all experiments, indel frequency was calculated as the number of discarded reads divided by the total number of reads ((number of indel-containing reads)/(number of reference-aligned reads). EditR (Kluesner et al., 2018) was used for PE activity quantifications from all sanger sequencing data.

### Live cell CRISPR-based DNA imaging

All live-cell imaging was carried out on a DeltaVision Ultra imaging system (GE Healthcare). The cells were cultured on No. 1.0 glass bottom dishes (MatTek). The microscope stage incubation chamber was maintained at 37°C and 5% CO2. GFP was excited at 488 nm and collected using filter at 498/30 nm (wavelength/bandwidth). Imaging data were acquired by DeltaVision Elite imaging (GE Healthcare Inc.) software. For the representative images, the raw data were deconvoluted by softWoRx (GE Healthcare Inc.) software.

### Off-target analysis

Potential off-target sites were predicted in the human genome (GRCh38/hg38) with Cas-OFFinder6 (http://www.rgenome.net/cas-offinder); The region around the off-target sites was amplified with Phanta Max Super-Fidelity DNA Polymerase (Vazyme), and subjected to high-throughput sequencing. The amplicons were analyzed with CRIPResso2 (V2.0.43) and the off-target sites are listed in Supplementary information. Primers used are listed in Supplementary information.

**Figure S1.**
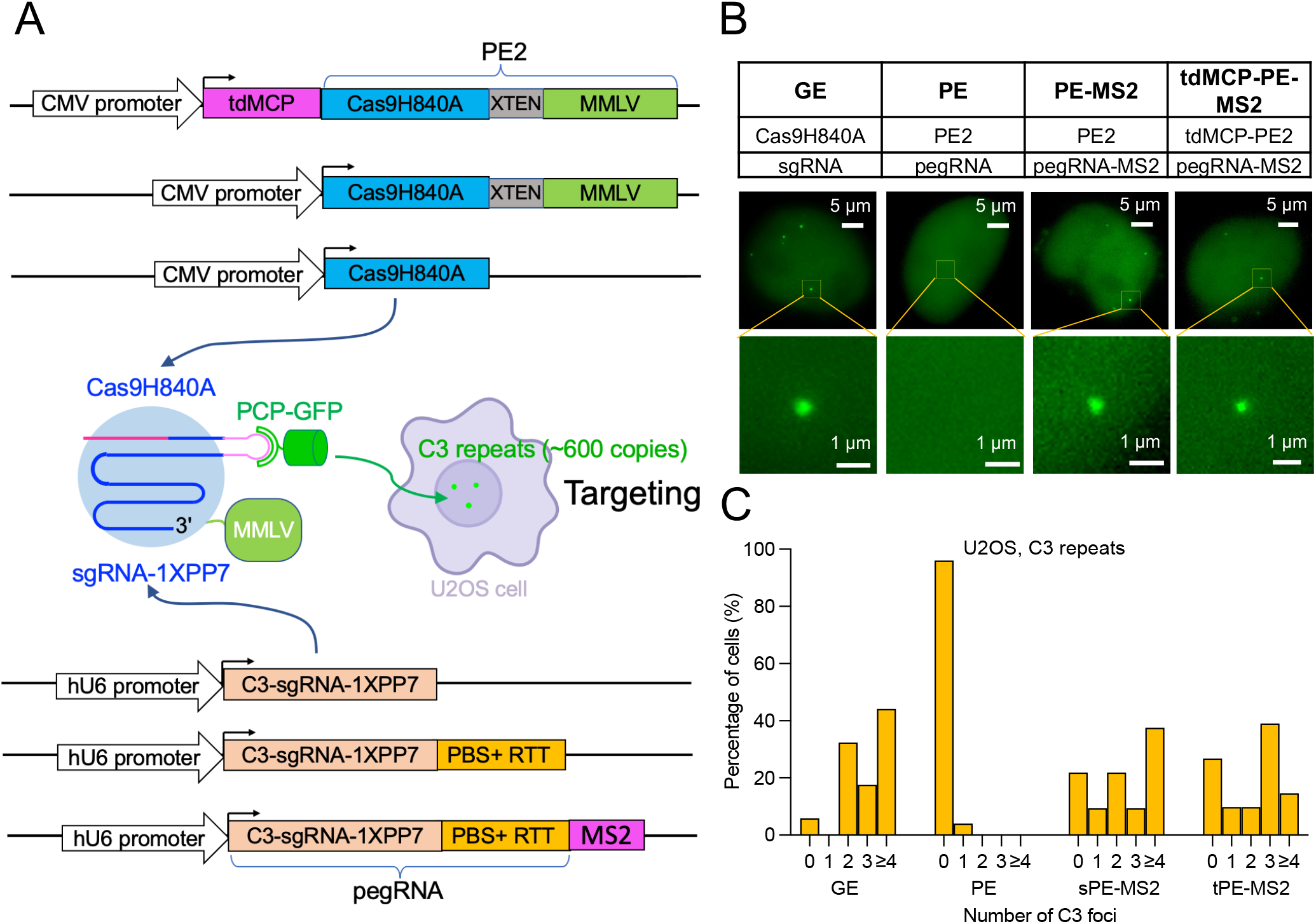
Increase of targeting efficiency by adding stem-loop RNA aptamers at the 3’-terminal of pegRNAs. (A) Schematic of the live-cell detection system for targeting efficiency. The detection system consists of Cas9 nickase, PCP-GFP, sgRNA-1XPP7 targeting C3 repeats (∼600 copies of target sites on chromosome 3). The expression of Cas9H840A, Cas9H840A-MMLV (PE2) or tdMCP-PE2 was driven by CMV promoter. The expression of sgRNA, pegRNA or pegRNA-MS2 was driven by human U6 (hU6) promoter. (B) U2OS cells were co-transfected PCP-GFP along with expression plasmids as indicated for GE, PE, sPE-MS2, or tPE-MS2. After 48 hours, cells were imaged by collecting z-stack images to capture all foci in each nucleus examined. Representative images show the 2D projection of 3D imaging. (C) Histograms showing the number of C3 foci per cell counts by CRISPR-based labeling, n=50 transfected cells of GE, PE, sPE-MS2 or tPE-MS2.

**Figure S2.**
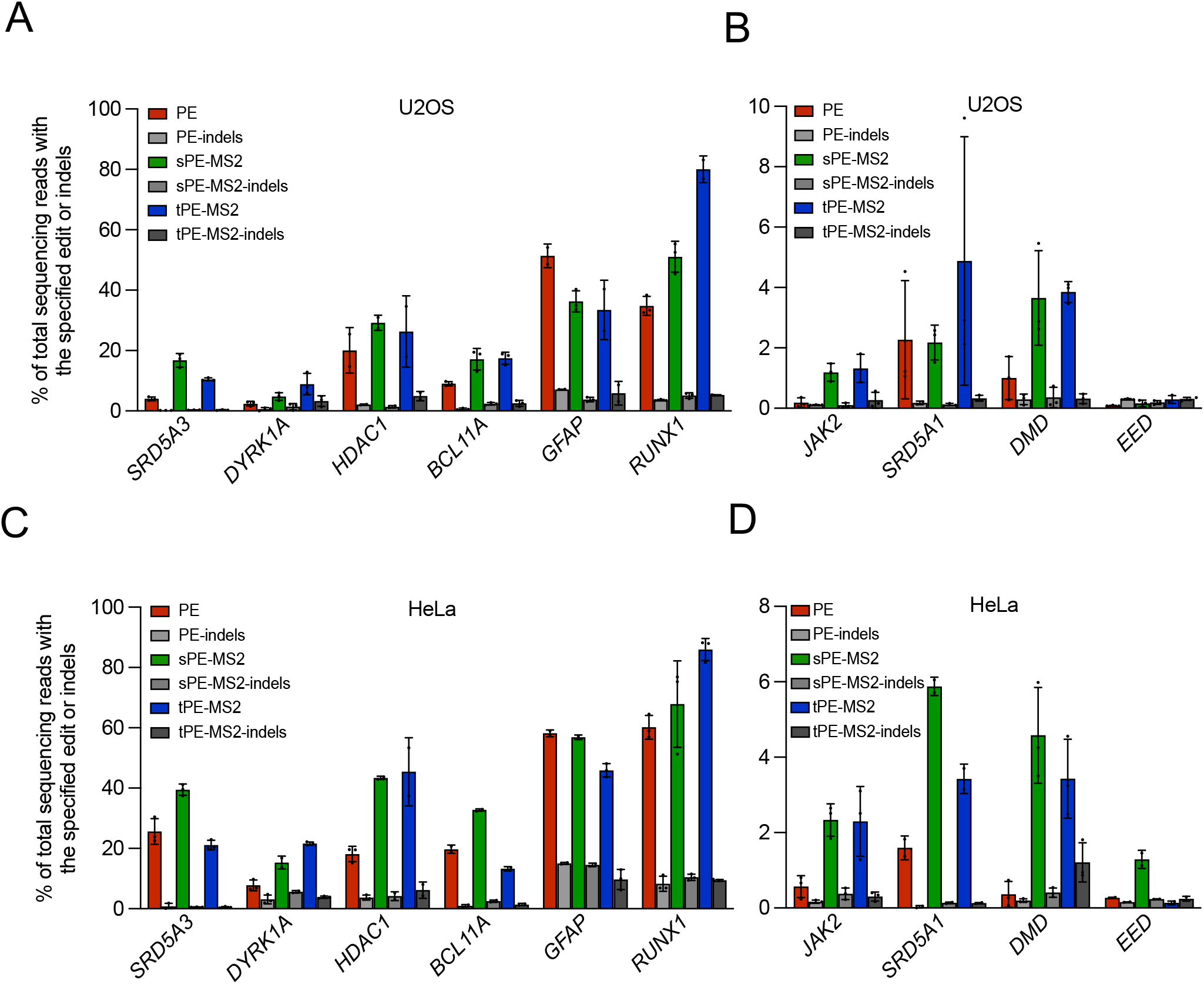
sPEs and tPEs increase PE efficiency at various loci in different cell lines. Efficiency of PE, sPE-MS2 and tPEb-MS2 mediated point mutation of *SRD5A3_*+2 C·G to A·T, *DYRK1A_*+1 C·G to G·C, *HDAC1_*+1 C·G to G·C, *BCL11A*_+1C·G to A·T, *GFAP*_+1A·T to T·A and *RUNX1*_+5 G·C to T·A using PE3 in U2OS cells (A), Hela cells (C). Efficiency of PE, sPE-MS2 and tPE-MS2 mediated point mutation of *DMD_*+1 T·A to C·G, *EED_*+1 A·T to T·A, *JAK2 _*+1 C·G to T·A and *SRD5A1*_+1 C·G to A·T using PE3 in U2OS cells (B), Hela cells (D). Data and error bars in (A-D) indicate the mean and standard deviation of three independent biological replicates.

**Figure S3.**
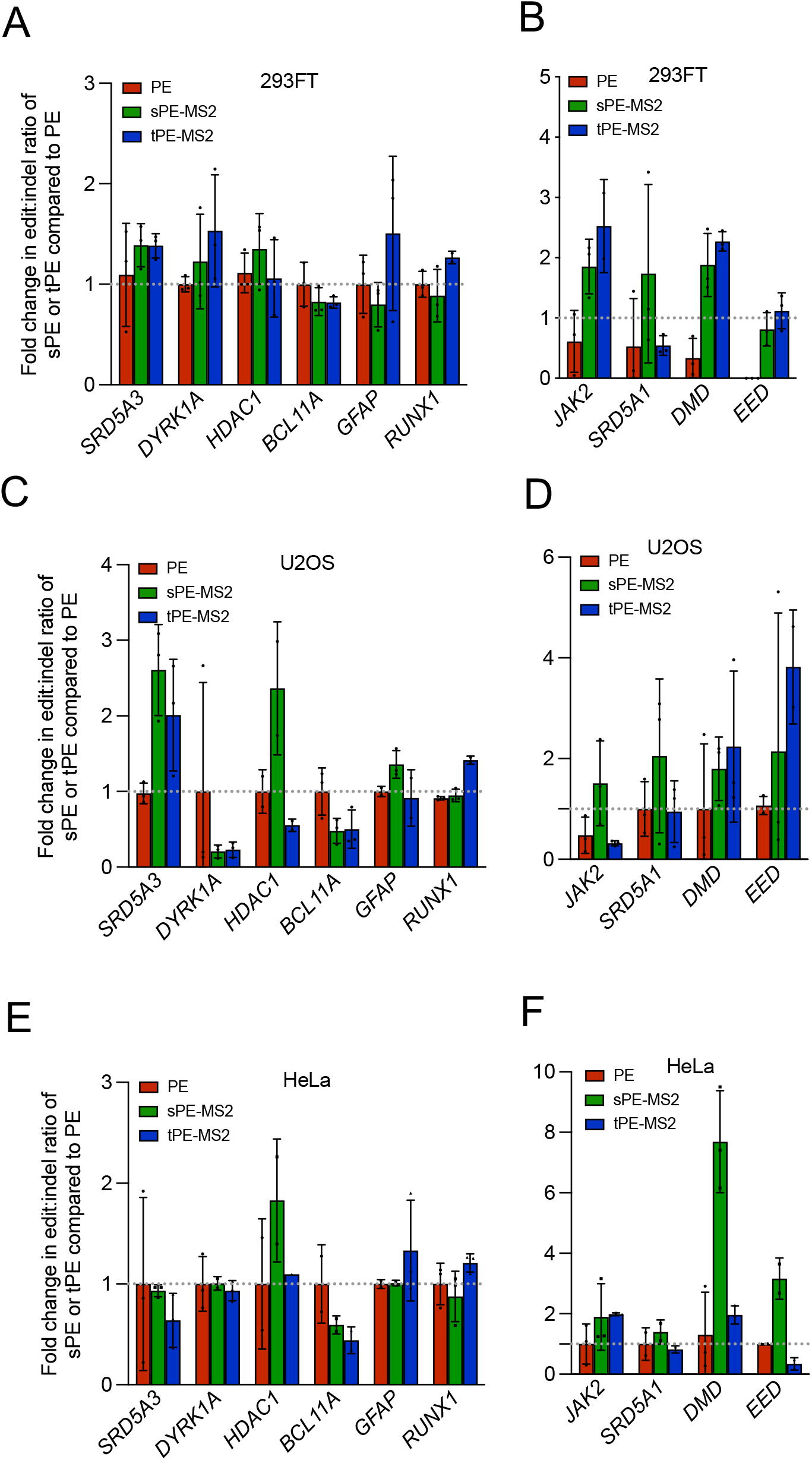
Fold change in correct editing and Edit:indel ratio for sPE-MS2 and tPE-MS2 at different loci shown in Figure 1 and Figure S2. Fold-change in the observed edit:indel ratio of PE, sPE-MS2 and tPE-MS2 mediated point mutation for *SRD5A3_*+2 C·G to A·T, *DYRK1A_*+1 C·G to G·C, *HDAC1_*+1 C·G to G·C, *BCL11A*_+1C·G to A·T, *GFAP*_+1A·T to T·A and *RUNX1*_+5 G·C to T·A in HEK293FT cells (A) U2OS cells (C) and HeLa cells (E). Fold-change in the observed edit:indel ratio of PE, sPE-MS2 and tPEb-MS2 mediated point mutation of *DMD_*+1 T·A to C·G, *EED_*+1 A·T to T·A, *JAK2_*+1 C·G to T·A and *SRD5A1*_+1 C·G to A·T in HEK293FT cells (B) U2OS cells (D) and HeLa cells (F). Values were calculated from the data presented in Figure 1 and Figure S2. Data and error bars indicate the mean and standard deviation of three in dependent biological replicates.

**Figure S4.**
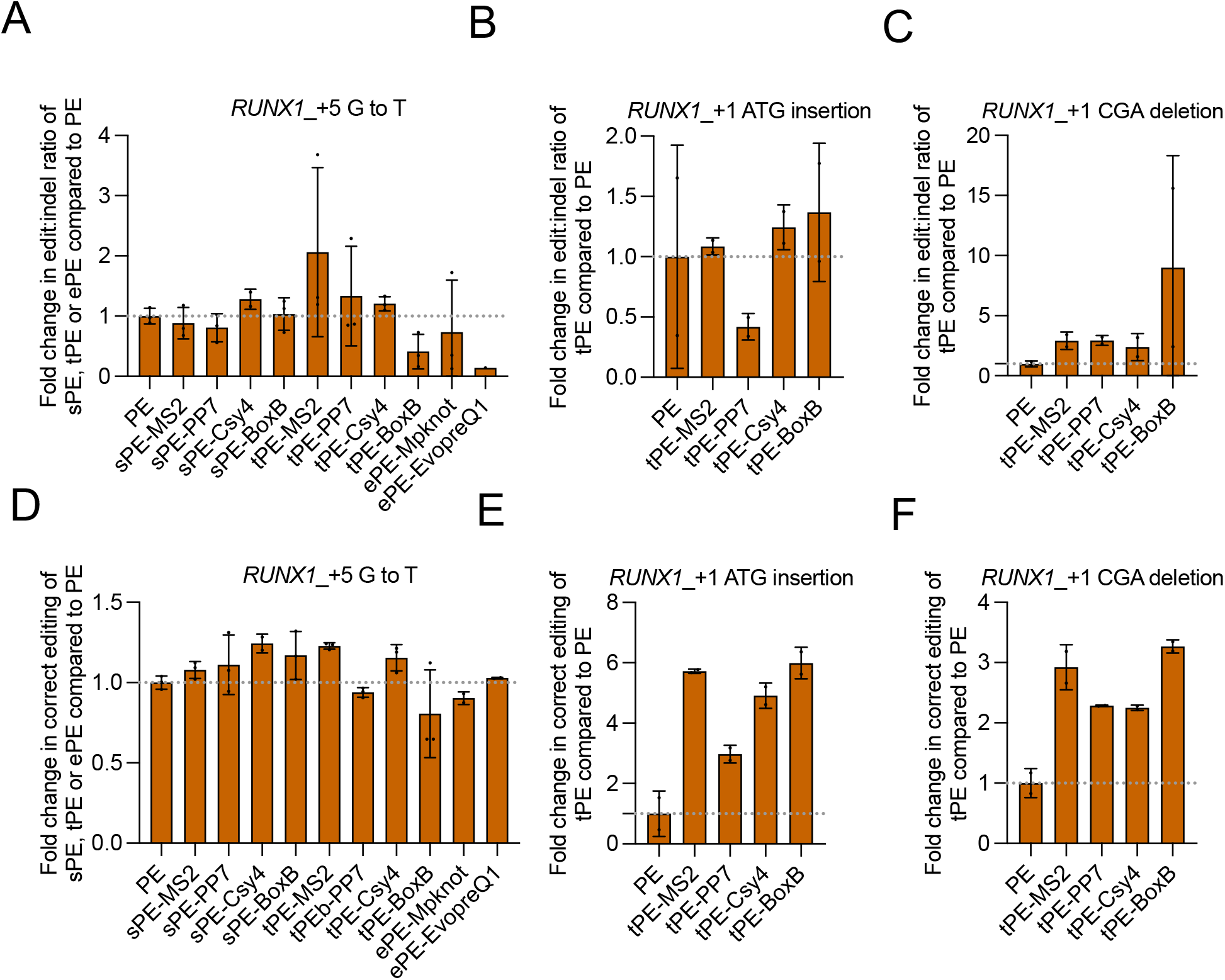
Fold change in correct editing and edit:indel ratio for sPEs and tPEs shown in Figure 1. (A) Fold-change in the observed edit:indel ratio of PE, sPE-MS2, tPE-MS2, ePE-Mpknot and ePE-EvopreQ1 mediated with point mutation for *RUNX1*_+5 G·C to T·A in HEK293FT cells. (B) Fold-change in the observed edit:indel ratio of PE, tPE-MS2, tPE-PP7, tPE-Csy4 and tPE-BoxB mediated with insertion for *RUNX1*_+1 ATG in HEK293FT cells. (C) Fold-change in the observed edit:indel ratio of PE, tPE-MS2, tPE-PP7, tPE-Csy4 and tPE-BoxB mediated with deletion for *RUNX1*_+1 CGA in HEK293FT cells. (D) Fold-change in the observed correct editing of PE, sPE-MS2, tPE-MS2, ePE-Mpknot and ePE-EvopreQ1 mediated with point mutation for *RUNX1*_+5 G·C to T·A in HEK293FT cells. (E) Fold-change in the observed correct editing of PE, tPE-MS2, tPE-PP7, tPE-Csy4 and tPE-BoxB mediated with insertion for *RUNX1*_+1 ATG in HEK293FT cells. (F) Fold-change in the observed correct editing of PE, tPE-MS2, tPE-PP7, tPE-Csy4 and tPE-BoxB mediated with deletion for *RUNX1*_+1 CGA in HEK293FT cells. Values were calculated from the data presented in Figure 1. Data and error bars indicate the mean and standard deviation of three in dependent biological replicates.

**Figure S5.**
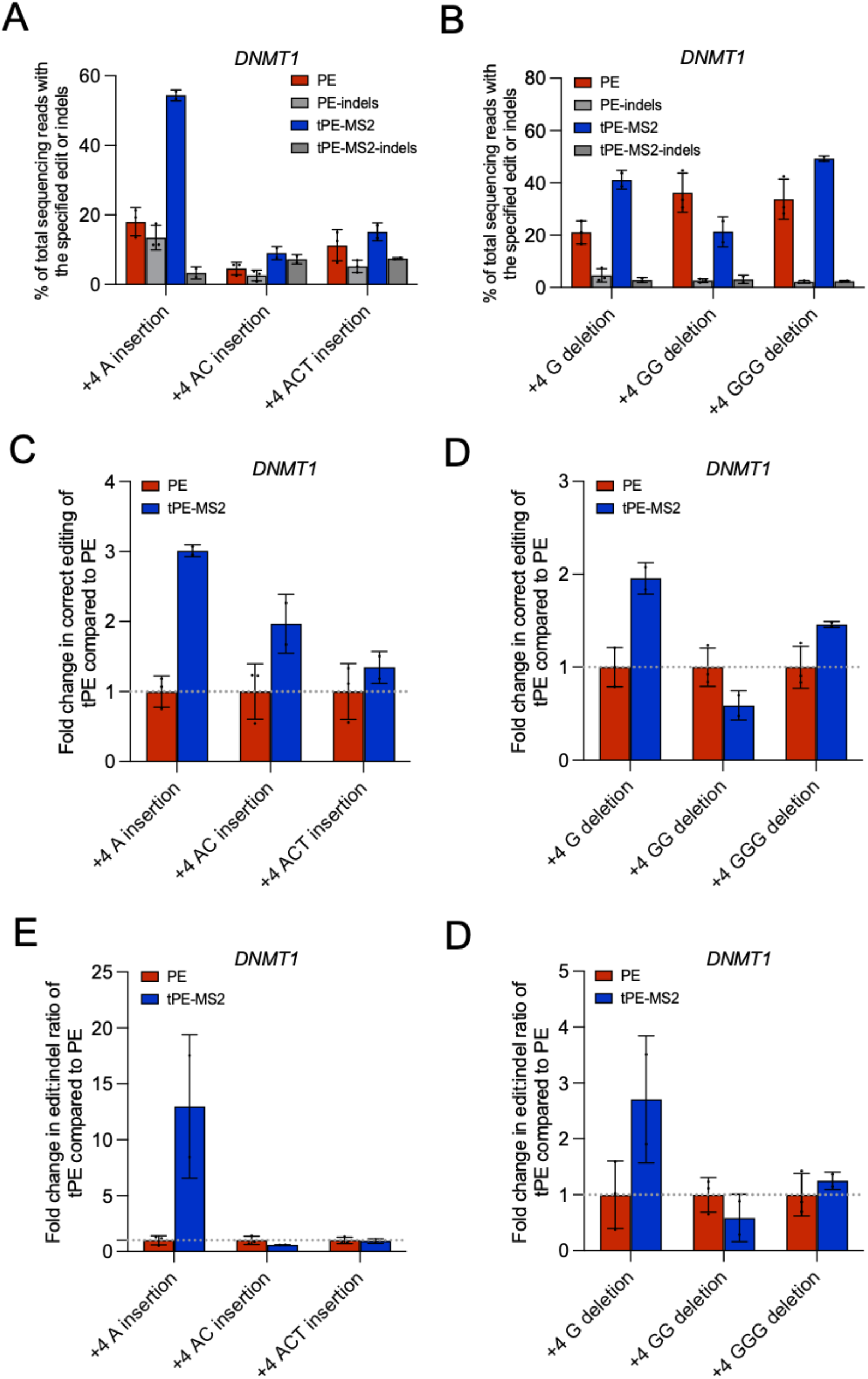
Insertion and deletion efficiency for *DNMT1* loci and fold change in correct editing and edit:indel ratio. (A) The efficiency of PE and tPE-MS2 mediated insertion of DNMT1_+4 A, +4 AC, +4 ACT using PE3 in 293FT cells. (B) The efficiency of PE and tPE-MS2 mediated deletion of DNMT1_+4 G, +4 GG, +4 GGG using PE3 in HEK293FT cells. (C) Fold-change in the observed correct editing ratio of PE, tPE-MS2 mediated with insertion for *DNMT1*_+4 A, *DNMT1_*+4 AC, *DNMT1_*+4ACT in HEK293FT cells. (D) Fold-change in the observed correct editing ratio of PE, tPE-MS2 mediated with deletion for *DNMT1*_+4 G, *DNMT1_*+4 GG, *DNMT1_*+4GGG in HEK293FT cells. (E) Fold-change in the observed edit:indel ratio of PE, tPE-MS2 mediated with insertion for *DNMT1*_+4 A, *DNMT1_*+4 AC, *DNMT1_*+4ACT in HEK293FT cells. (F) Fold-change in the observed edit:indel ratio of PE, tPE-MS2 mediated with deletion for *DNMT1*_+4 G, *DNMT1_*+4 GG, *DNMT1_*+4GGG in HEK293FT cells. Values were calculated from the data presented in Figure S5A and Figure S5B. Data and error bars indicate the mean and standard deviation of three in dependent biological replicates.

**Figure S6.**
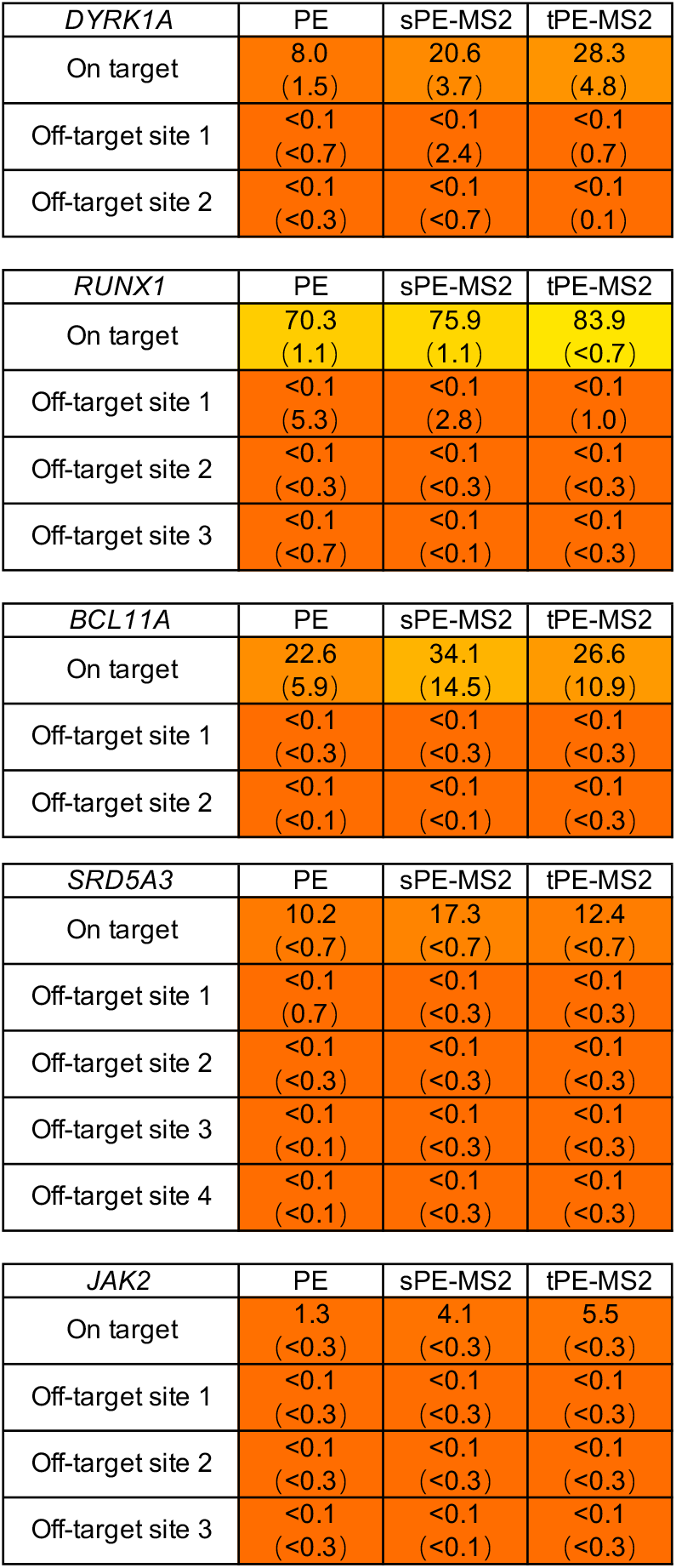
Comparison of off-target activity between canonical PE, sPE-MS2 and tPE-MS2. on-target and off-target activities of PE, sPE-MS2 and tPE-MS2 at *DYRK1A, RUNX1, BCL11A, SRD5A3* and *JAK2* were detected. PE, sPE and tPE editing is shown as % prime editing alongside % indels (in parentheses). Circular prime RNA (cpRNA)

**Figure S7.**
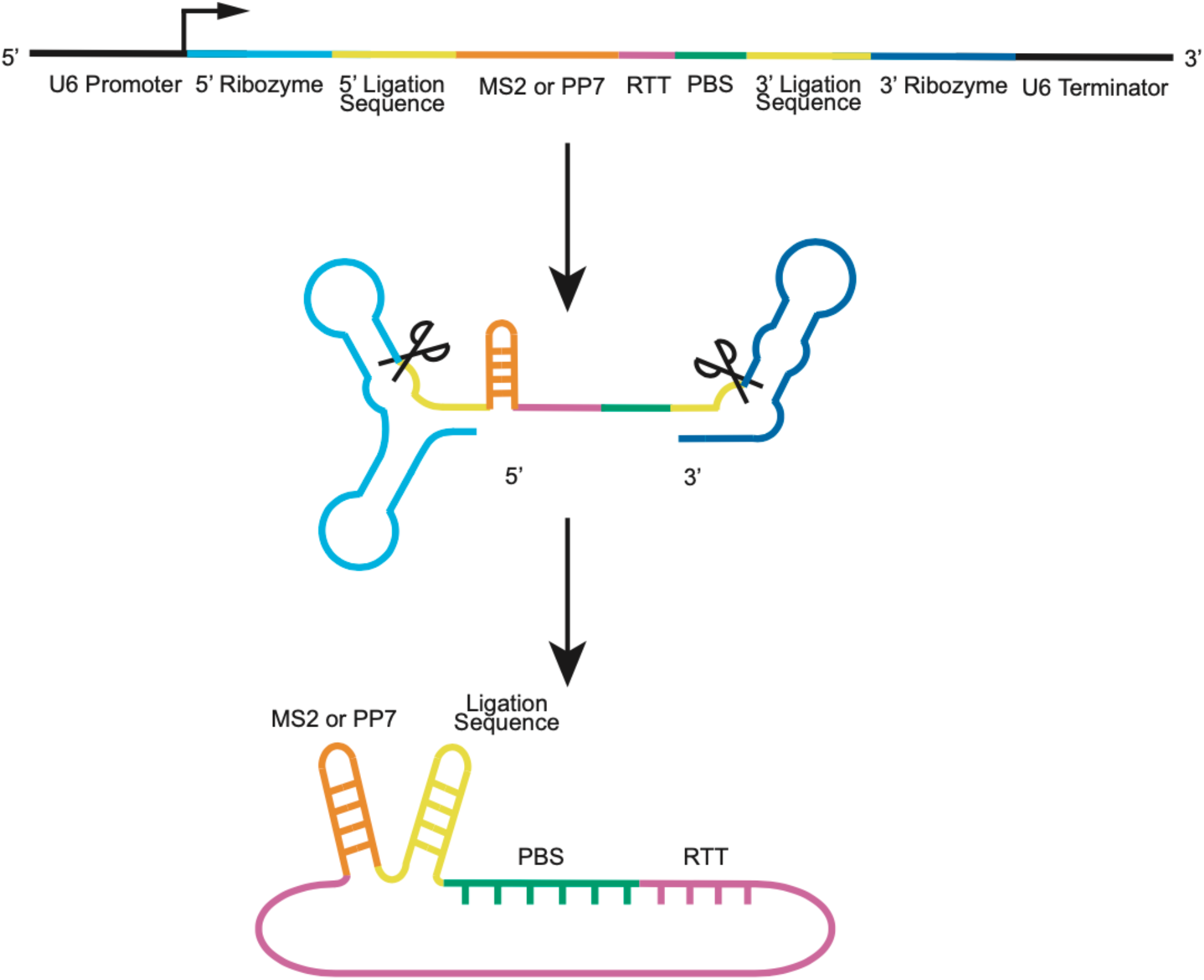
Circular prime RNA generated by the Tornado expression system used in Figure 2. Construct design for generating autocatalytically processed circular prime RNA (cpRNA) by the Tornado expression system. The circular prime RNA in the Tornado cassette is driven by U6 promoter. The circular pRNA consists of MS2 or PP7 (orange), RTT (reddish-purple) and PBS (bluish-green). The circular pRNA is flanked by the 5′- and 3′-stem-forming sequences (yellow), each of which is flanked by the 5′- and 3′-self-cleaving ribozymes (sky blue and blue, respectively). The stem-forming by 5’- and 3’-ligation sequences facilitates the circularization of pRNA into circular prime RNA (cpRNA).

**Figure S8.**
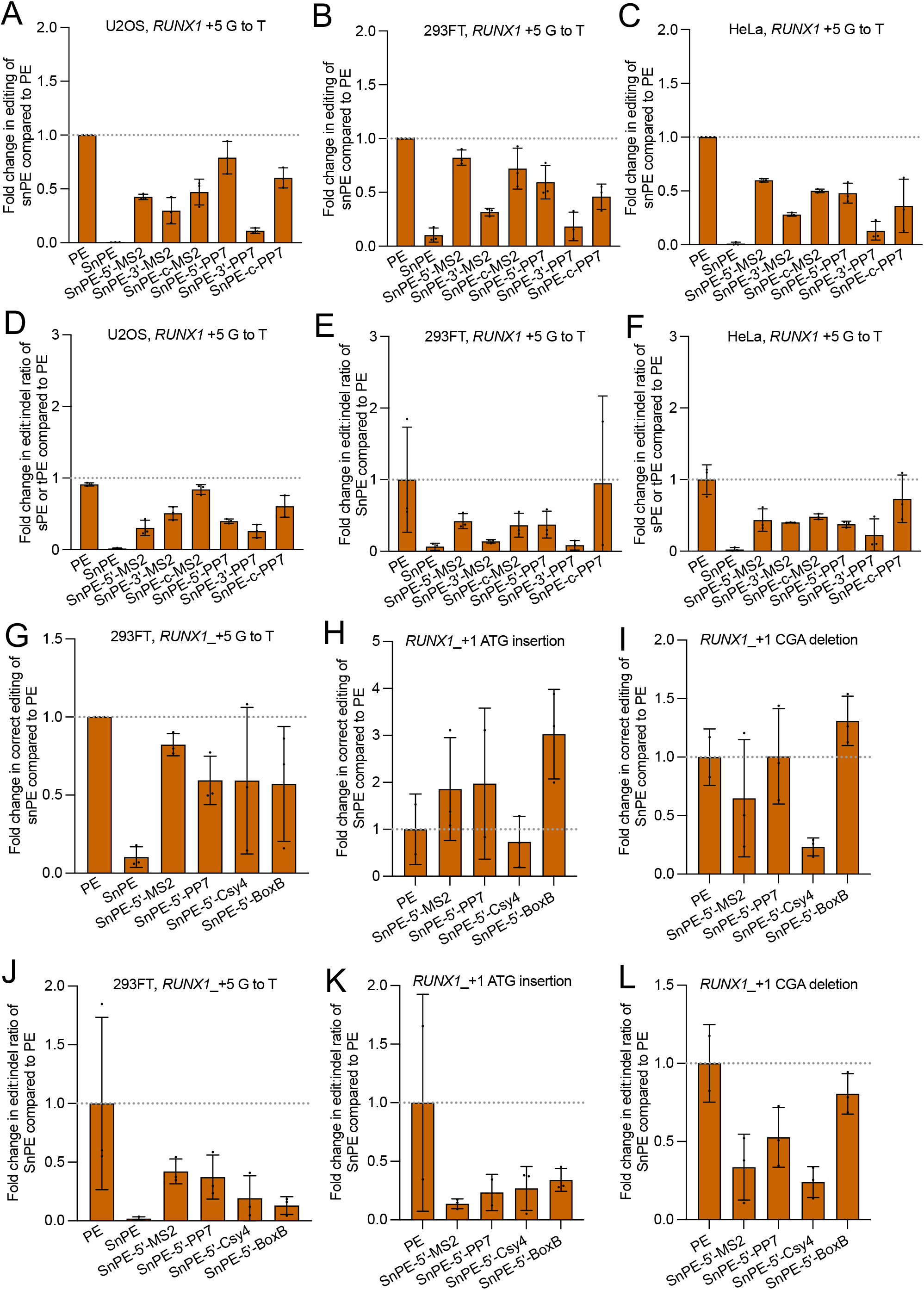
Fold change in correct editing and edit:indel ratio for SnPEs shown in Figure 2. Fold-change in the observed correct editing for *RUNX1*_+5 G·C to T·A transversion of SnPE-5’-MS2 or PP7, SnPE-3’-MS2 or PP7, SnPE-c-MS2 or PP7, compared to canonical PE (dashed line) in U2OS cells (A), HEK293FT cells (B), and HeLa cells (C). Fold-change in the observed edit:indel ratio for *RUNX1*_+5 G·C to T·A of SnPE-5’-MS2 or PP7, SnPE-3’-MS2 or PP7, SnPE-c-MS2 or PP7, compared to canonical PE (dashed line) in U2OS cells (D), HEK293FT cells (E), and HeLa cells (F). (G) Fold-change in the observed correct edting for SnPE-5’-MS2, SnPE-5’-PP7, SnPE-5’-Csy4 and SnPE-5’-boxB mediated with *RUNX1*_+5 G·C to T·A in HEK293FT cells. (H) Fold-change in the observed correct editing of PE, tPE-MS2, tPE-PP7, tPE-Csy4 and tPE-BoxB mediated with insertion for *RUNX1*_+1 ATG in HEK293FT cells. (I) Fold-change in the observed correct editing of PE, tPE-MS2, tPE-PP7, tPE-Csy4 and tPE-BoxB mediated with deletion for *RUNX1*_+1 CGA in HEK293FT cells. (J) Fold-change in the observed edit:indel ratio for SnPE-5’-MS2, SnPE-5’-PP7, SnPE-5’-Csy4 and SnPE-5’-boxB mediated with *RUNX1*_+5 G·C to T·A in HEK293FT cells. (K) Fold-change in the observed edit:indel ratio of PE, tPE-MS2, tPE-PP7, tPE-Csy4 and tPE-BoxB mediated with insertion for *RUNX1*_+1 ATG in HEK293FT cells. (L) Fold-change in the observed edit:indel ratio of PE, tPE-MS2, tPE-PP7, tPE-Csy4 and tPE-BoxB mediated with deletion for *RUNX1*_+1 CGA in HEK293FT cells. Values were calculated from the data presented in Figure 2. Data and error bars reflect the mean and standard deviation of three independent biological replicates.

**Figure S9.**
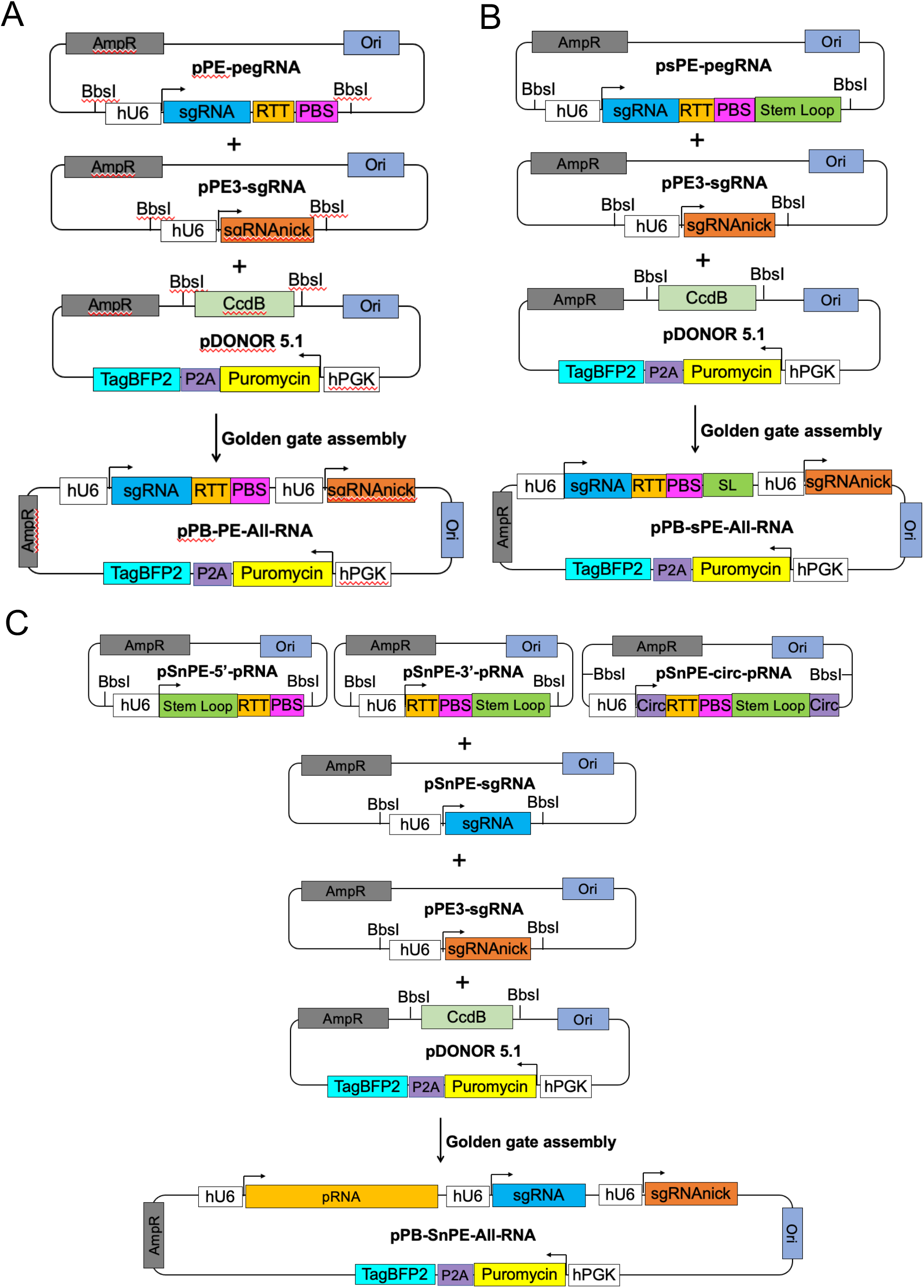
Cloning strategy for single construct containing all RNA expression components for modified or split pegRNA prime editors. Construction of pPB-PE-ALL-RNA for canonical PE, pPB-sPE-ALL-RNA plasmids for sPEs and tPEs, and pPB-SnPE-ALL-RNA for SnPEs. (A) pPB-PE-ALL-RNA was reconstituted from pPE-pegRNA, pPE3-sgRNA and pDONOR5.1 by golden gate assembly, pPE-pegRNA consists of sgRNA (blue), RTT (gold) and PBS (Magenta), pPE3-sgRNA contains sgRNAnick (orange), pDONOR5.1 consists of CcdB gene (green), Puromycin (yellow), P2A (purple) and TagBFP2 (Cyan). (B) pPB-sPE-ALL-RNA was reconstituted from psPE-pegRNA, which contains sgRNA (blue), RTT (gold), PBS (Magenta) and stem-loop RNA aptamer (dark green), pPE3-sgRNA and pDONOR5.1 were the same in (A). (C) pPB-SnPE-ALL-RNA contains three types of pRNA: pSnPE-5’-pRNA, pSnPE-3’-pRNA and pSnPE-circ-pRNA. pSnPE-5’-pRNA consists of stem loop at 5’-terminal (dark green), RTT (gold) and PBS (Magenta). pSnPE-3’-pRNA contains stem-loop at 3’-terminal (dark green), RTT (gold) and PBS (Magenta). pSnPE-circ-pRNA contains circular sequence located at both terminals (purple), stem loop (dark green), RTT (gold) and PBS (Magenta). pSnPE-sgRNA was pSnPE-pRNAs, sgRNA (blue), pPE3-sgRNA and pDONOR5.1 that were used in (A) and (B).

## Notes

### Competing Interest Statement

The authors have declared no competing interest.

## References

Anzalone AV, Gao XD, Podracky CJ, Nelson AT, Koblan LW, Raguram A, Levy JM, Mercer JAM, Liu DR (2021) Programmable deletion, replacement, integration and inversion of large DNA sequences with twin prime editing. Nat. Biotechnol., doi: 10.1038/s41587-021-01133-w

Anzalone AV, Randolph PB, Davis JR, Sousa AA, Koblan LW, Levy JM, Chen PJ, Wilson C, Newby GA, Raguram A et al. (2019) Search-and-replace genome editing without double-strand breaks or donor DNA. Nature 576:149–157.

Chen PJ, Hussmann JA, Yan J, Knipping F, Ravisankar P, Chen PF, Chen C, Nelson JW, Newby GA, Sahin M. et al. (2021) Enhanced prime editing systems by manipulating cellular determinants of editing outcomes. Cell 184:5635–5652.

Choi J, Chen W, Suiter CC, Lee C, Chardon FM, Yang W, Leith A, Daza RM, Martin B, Shendure J. (2021) Precise genomic deletions using paired prime editing. Nat. Biotechnol. 40:218–226.

Cong L, Ran FA, Cox D, Lin S, Barretto R, Habib N, Hsu PD, Wu X, Jiang W, Marraffini LA et al. (2013) Multiplex genome engineering using CRISPR/Cas systems. Science 339:819–823.

Convery MA, Rowsell S, Stonehouse NJ, Ellington AD, Hirao I, Murray JB, Peabody DS, Phillips SE, Stockley PG. (1998) Crystal structure of an RNA aptamer-protein complex at 2.8 A resolution. Nat. Struct Biol. 5:133–139.

Gaudelli NM, Komor AC, Rees HA, Packer MS, Badran AH, Bryson DI, Liu DR. (2017) Programmable base editing of A•T to G•C in genomic DNA without DNA cleavage. Nature 551:464–471.

Haurwitz RE, Jinek M, Wiedenheft B, Zhou K, Doudna JA. (2010) Sequence-and structure-specific RNA processing by a CRISPR endonuclease. Science, 329:1355–1358.

Hendel A, Bak RO, Clark JT, Kennedy AB, Ryan DE, Roy S, Steinfeld I, Lunstad BD, Kaiser RJ, Wilkens AB. et al. (2015) Chemically modified guide RNAs enhance CRISPR-Cas genome editing in human primary cells. Nat. Biotechnol., 33:985–989.

Jiang T, Zhang XO, Weng Z, Xue W. (2021) Deletion and replacement of long genomic sequences using prime editing. Nat. Biotechnol., doi: 10.1038/s41587-021-01026-y

Koblan LW, Arbab M, Shen MW, Hussmann JA, Anzalone AV, Doman JL, Newby GA, Yang D, Mok B, Replogle JM et al. (2021) Efficient C•G-to-G•C base editors developed using CRISPRi screens, target-library analysis, and machine learning. Nat. Biotechnol. 39:1414–1425.

Komor AC, Kim YB, Packer MS, Zuris JA, Liu DR. (2016) Programmable editing of a target base in genomic DNA without double-stranded DNA cleavage. Nature 533:420–424.

Kosicki M, Tomberg K, Bradley A. (2018) Repair of double-strand breaks induced by CRISPR–Cas9 leads to large deletions and complex rearrangements. Nat. Biotechnol. 36:765–771.

Kurt IC, Zhou R, Iyer S, Garcia SP, Miller BR, Langner LM, Grünewald J, Joung JK. (2021) CRISPR C-to-G base editors for inducing targeted DNA transversions in human cells. Nat. Biotechnol. 39:41–46.

Landrum MJ, Lee JM, Benson M, Brown G, Chao C, Chitipiralla S, Gu B, Hart J, Hoffman D, Hoover J. et al. (2016) ClinVar: public archive of interpretations of clinically relevant variants. Nucleic Acids Res. 44:D862–D868.

Lin Q, Zong Y, Xue C, Wang S, Jin S, Zhu Z, Wang Y, Anzalone AV, Raguram A, Doman JL et al. (2020) Prime genome editing in rice and wheat. Nat. Biotechnol. 38:582–585.

Litke JL, Jaffrey SR. (2019) Highly efficient expression of circular RNA aptamers in cells using autocatalytic transcripts. Nat. Biotechnol. 37:667–675.

Liu Y, Yang G, Huang S, Li X, Wang X, Li G, Chi T, Chen Y, Huang X, Wang X. (2021) Enhancing prime editing by Csy4-mediated processing of pegRNA. Cell Res. 31:1134–1136.

Ma H, Tu LC, Naseri A, Huisman M, Zhang S, Grunwald D, Pederson T. (2016a) CRISPR-Cas9 nuclear dynamics and target recognition in living cells. J. Cell Biol. 214:529–537.

Ma H, Tu LC, Naseri A, Chung YC, Grunwald D, Zhang S, Pederson T. (2018) CRISPR-Sirius: RNA scaffolds for signal amplification in genome imaging. Nat. Methods. 15:928–931.

Ma H, Tu LC, Naseri A, Huisman M, Zhang S, Grunwald D, Pederson T. (2016b) Multiplexed labeling of genomic loci with dCas9 and engineered sgRNAs using CRISPRainbow. Nat. Biotechnol. 34:528–530.

Mali P, Yang L, Esvelt KM, Aach J, Guell M, DiCarlo JE, Norville JE, Church GM. (2013) RNA-guided human genome engineering via Cas9. Science 339:823–826.

Nelson JW, Randolph PB, Shen SP, Everette KA, Chen PJ, Anzalone AV, An M, Newby GA, Chen JC, Hsu A et al. (2021) Engineered pegRNAs improve prime editing efficiency. Nat. Biotechnol. 40:402–410.

Bae S, Park J, Kim JS. (2014) Cas-OFFinder: a fast and versatile algorithm that searches for potential off-target sites of Cas9 RNA-guided endonucleases, Bioinformatics, 30: 1473–1475.

Stanton BZ, Chory EJ, Crabtree GR. (2018) Chemically induced proximity in biology and medicine. Science, 359:eaao5902.

Urbanek MO, Galka-Marciniak P, Olejniczak M, Krzyzosiak WJ. (2014) RNA imaging in living cells - methods and applications. RNA Biol. 11:1083–1095.

Wu Y, Liu Y, Huang Z, Wang X, Jin Z, Li J, Limsakul P, Zhu L, Allen M, Pan Y et al. (2021) Control of the activity of CAR-T cells within tumours via focused ultrasound. Nat. Biomed Eng. 5:1336–1347.

Yu Y, Wu X, Guan N, Shao J, Li H, Chen Y, Ping Y, Li D, Ye H. (2020) Engineering a far-red light-activated split-Cas9 system for remote-controlled genome editing of internal organs and tumors. Sci. Adv. 6:eabb1777.

Yin H, Song CQ, Suresh S, Wu Q, Walsh S, Rhym LH, Mintzer E, Bolukbasi MF, Zhu LJ, Kauffman K et al. (2017) Structure-guided chemical modification of guide RNA enables potent non-viral in vivo genome editing. Nat. Biotechnol., 35:1179–1187.

Zalatan JG, Lee ME, Almeida R, Gilbert LA, Whitehead EH, La Russa M, Tsai JC, Weissman JS, Dueber JE, Qi LS et al. (2015) Engineering complex synthetic transcriptional programs with CRISPR RNA scaffolds. Cell 160:339–350.

Zhao D, Li J, Li S, Xin X, Hu M, Price MA, Rosser SJ, Bi C, Zhang X. (2021) Glycosylase base editors enable C-to-A and C-to-G base changes. Nat. Biotechnol. 39:35–40.

Zhou Y, Kong D, Wang X, Yu G, Wu X, Guan N, Weber W, Ye H. (2021) A small and highly sensitive red/far-red optogenetic switch for applications in mammals. Nat. Biotechnol. 40:262–272.

